# Indigenous communities’ perceptions of ecosystem services of a montane forest ecosystem in Nigeria: Implications for restoration planning

**DOI:** 10.64898/2026.10.28.762469

**Authors:** Iveren Abiem, Ahmed Difa Ali, Emmanuel Barde Elisha, Gabriel Atsen Dabo, Christopher Tumba Yohanna, Susinya Habila, Danladi Mohammed Umar, Hazel Chapman

## Abstract

Restoration of protected areas should aim to maximize outcomes for both biological conservation and local people’s livelihoods. In rural Africa, livelihoods are inextricably linked to the provision of ecosystem services (ES) from natural resources, so that community-centred biodiversity conservation must be guided by local perceptions of ES. Our study examined perceptions of ES among eight communities within a 5-km radius of a Nigerian forest reserve characterized by an elevational gradient of 700 m, spanning montane forest, mid-elevation forest, and savannah. The montane forest is noted for its outstanding biodiversity value. We employed semi-structured interviews, focus group discussions and species-use inventories to characterize which ES are recognised, and their perceived importance. We also conducted an ethnobotanical assessment to document community uses of plants and classified the species as savannah or forest species. Across settlements, we observed a three-fold variation in the number of ES recognized, ranging from 6 to 20. Provisioning services, especially food, were ranked higher than regulating, cultural, or supporting services in most settlements. Notably, most provisioning services were from savannah, rather than forest species. Our study adds to growing data on indigenous knowledge about ecosystem services. Implications from these findings are that: i) Landscape-scale management should maintain a mosaic of montane forest and derived savannah to reflect prevailing provisioning needs while enhancing regulating and supporting services. ii) Restoration should prioritise tangible provisioning benefits (e.g., non-timber forest products, medicines and food) alongside regulating and supporting services to align with local priorities. iii) Given potential trade-offs between scientifically derived valuations (biodiversity, regulation) and stakeholder priorities, co-design of restoration objectives and adaptive governance are essential to secure local buy-in and on-going stewardship.

## INTRODUCTION

Restoration of protected areas should maximize outcomes for both biological conservation and local people’s livelihoods. Protected areas safeguard biodiversity (De Fries et al., 2007), but at the same time support human welfare through the provision of ecosystem services (ES) (Costanza et al., 1997; Costanza et al., 2017). Ecosystem services are classified into provisioning, regulating, cultural, and supporting (Millennium Ecosystem Assessment, 2005; Mitsch & Gosselink, 2015). The ES framework is widely applied in conservation science and policy decision making; by connecting ecological processes to human wellbeing, the framework is used to guide protected area management, including their restoration (Shimamoto et al., 2017; Brouwer et al., 2024). For example, in Africa, active forest restoration might prioritize the planting of tree species which provide the benefits of high carbon sequestration (regulating), food and medicine (provisioning) as well as the maintenance of biodiversity (supporting). However, this ES approach can lead to tensions, especially in tropical areas with high biodiversity and high dependency on this diversity by local communities (Fashing et al., 2004; Camacho-Benavides et al., 2013). Targeting multiple ES introduces the potential for trade-offs (Beckline & Ntoko, 2022). Trade-offs occur when the increase in one ES leads to the reduction of another (Raudsepp-Hearne et al., 2010); for example, converting grasslands into forest may increase carbon sequestration at the expense of historical grassland biomes (Veldman et al., 2017).

Evaluating ES for protected area restoration requires an integrative approach that combines ecological, economic and socio-cultural criteria (Villegas-Palacio et al., 2016; Zhao et al., 2024; Tasi et al., 2025). The socio-cultural evaluation, which has been frequently overlooked (Tengö et al., 2017; Cuni-Sanchez et al., 2019) is an especially critical component, as it uses stakeholder perceptions to assess the value of ES (Martín-López et al., 2012), leading to more effective conservation management and restoration outcomes (Iniesta-Arandia et al., 2014; Kovács et al., 2015; Scholte et al., 2015; Velasco-Muñoz et al., 2022). Socio-cultural evaluations use data from stakeholders to infer the value of ecosystem (Martín-López et al., 2012). By integrating local values and perspectives, the socio-cultural approach not only complements, but also enriches purely ecological assessments. Ultimately, it ensures that management interventions are appropriately embedded within, and supported by, the local socio-cultural context.

As seen above, a single ecosystem often generates multiple ES, so that the communities dependent on that landscape may value it for diverse reasons, with significant variation in individual preferences (Miller et al., 2021). Consequently, it is important to understand how distinct community subgroups, such as individual settlements or specific ethnic groups, perceive and prioritize these services. By doing this, management strategies can be designed that are more likely to be accepted by the whole community (Bennett, 2015).

Africa’s highland regions are crucial for local livelihoods and global environmental health (Platts et al., 2011; Abera & Dagnew, 2025). These regions are also known to support the densest rural human populations in Africa which has resulted in intense pressure on forests, mostly due to overexploitation, agricultural expansion and habitat loss (Cronin et al., 2014). To counter this, policy and management of African highland protected areas is increasingly guided by outcomes from studies of local stakeholders’ perception of the ES montane environments provide (Osewe et al., 2024). However, such studies are overwhelmingly from East African highlands with a focus on forests. Studies from West African highlands are rare (but see Cuni-Sanchez et al., 2019) and none that we are aware of include a steep ecological transition from highland to mid-elevation, nor forest-savannah mosaics. None to our knowledge explicitly aim to inform forest restoration management.

This study sought to incorporate the perceptions of local stakeholders regarding the ES provided by a Nigerian montane/mid-altitude forest reserve located on the Mambilla Plateau on the Nigerian side of the Nigeria/Cameroon border. The reserve, a mosaic of high diversity forest within a degraded grassland/savannah matrix is representative of remnant Cameroon highland forests more generally. The forest is rich in Afromontane endemics and globally recognised for its biodiversity (Chapman et al., 2004), but the reserve is also important to local stakeholders-subsistence farmers and Fulani herdsmen, for the ES it provides. Gazetted as a Forest Reserve in 1969 the law allows for the collection of certain fruits and thatching grass. Cattle grazing and hunting of bush meat is illegal. However, as with most of these forests, it is threatened by fire, cattle grazing and farmland encroachment (Chapman et al., 2004; Cheek et al., 2000). By understanding how stakeholders perceive the ES provided by the reserve, we aimed to maximise restoration value for communities whose livelihoods depend, to varying degrees, on this montane ecosystem. In doing so, this work contributes to the Nigerian Montane Forest Project (NMFP), which seeks to optimize the spatial configuration and species composition of restoration plantings to enhance ES (Loan et al., 2025) and to inform a broader restoration strategy for protected areas along the 1800 km Cameroon Volcanic Line (Koch et al., 2012). Specifically, we asked:

1. How do eight settlements within 5km of the Ngel Nyaki Forest Reserve perceive the ES provided by the reserve?
2. How are the perceived ES divided between the highly diverse Afromontane/submontane forest with high conservation value and the degraded mid altitude forest and derived savanna /grassland?
3. How will our findings inform restoration strategies?

## METHODS

### Study Area

As part of the Cameroon highlands ecoregion (Dolinay et al., 2021), the Mambilla plateau in Taraba State, Nigeria, lies within the ‘Guinean Forests Hotspot’ centre of endemism (Cronin et al., 2014). The highlands are well documented for floristic, avian, amphibian and mammalian diversity (Cheek et al. 2000; Chapman et al. 2004; Nsor et al., 2019; Abiem et al., 2020; BakwoFils et al., 2021; Dolinay et al., 2021; Tela et al., 2021;).

Our surveys targeted settlements on the western escarpment of the plateau, all within 5 km of the Ngel Nyaki Forest Reserve, close to the Cameroonian border. The reserve is 4,600 ha, extending between 07°05’ N latitude and 11°05’ E longitude, with an elevational gradient from 900 to 1,600 m in elevation. Mean annual rainfall is ∼1,800 mm (mainly April–October) and mean annual plateau temperature is ∼19°C (Lai et al., 2025). The vegetation gradient covers montane/submontane to mid/lowland forest and associated grassland and derived savannah. The montane forest and grasslands host Cameroon Highland endemics and Red Data-listed taxa (Chapman & Chapman 2001; Borokini et al., 2012; Blackburn, 2010; Osinubi et al., 2018). The area is recognised as a Key Biodiversity Area (Key Biodiversity Areas partnership, 2026).

At the upper limits of forest (∼ 1500-1600 elevation), common tree species include *Strombosia schefflera, Macaranga occidentalis, Carapa oreophila*, *Eugenia gilgii*, *Pleiocarpa pycnantha, Pouteria altissima* and *Symphonia globulifera.* The grasslands within the reserve are dominated by *Hyparrhenia* species with scattered trees such as *Dombeya ledermannii, Harungana madagascariensis, Croton macrostachyus* and *Bridelia speciosa.* Lower slopes and valleys (∼900-m) support forest/savannah with sub-montane-lowland affinities such as *Terminalia leiocarpa, Vitex doniana*, *Afzelia africana* and *Pterocarpus erinaceus* (Abiem et al 2020; Chapman and Chapman 2001).

Despite being assigned a State Forest Reserve status in 1969, Ngel Nyaki faces ongoing pressures from farming and Fulani pastoralism. Burning and grazing have created derived savannah across much of the reserve (Chapman et al. 2004;). Between 2008–2011, the Nigerian Conservation Foundation and BirdLife International conducted stakeholder surveys and workshops around Ngel Nyaki forest reserve to promote sustainable land use. While participation issues limited impact, beekeepers benefited (Nigerian Conservation Foundation/Royal Society for Protection of Birds, 2011) and ideas of forest conservation were introduced to the communities who have not forgotten. Since then, the Nigerian Montane Forest Project (NMFP), in collaboration with the Taraba State Ministry of the Environment and Climate change have improved patrolling and fire/grazing protection, but the area remains highly vulnerable to anthropogenic pressures.

### Settlements surrounding Ngel Nyaki Forest Reserve

We sampled eight settlements located within 5 km of the Ngel Nyaki Forest reserve boundary; *Dombo Gishi, Dujere, Gidan Musa, Mai Wuya, Panso, Mayo Yembe, Yelwa* and *Zongo Ajiya* (Fig 1; Table 1). Settlement populations ranged in size from approximately 50 individuals (7 households; Panso) to over 2500 individuals (216 households; Yelwa). These communities comprise five primary ethnic groups: Mambilla, Kaka, Ndola, Kambu and Fulani (Elisha, 2023). The Mambilla, Kaka, Ndola, and Kambu are predominantly subsistence farmers and traders who share intertwined histories associated with the Mambilla Plateau and the contiguous Cameroon borderlands (Lenshie et al., 2023). In contrast the Fulani (also known as Fula or Fulbe) are a diverse group of traditionally semi-nomadic pastoralists, who have increasingly transitioned to a more sedentary lifestyle. Levels of education are low, only two of the eight settlements in our study have a secondary school.

**FIGURE 1:**
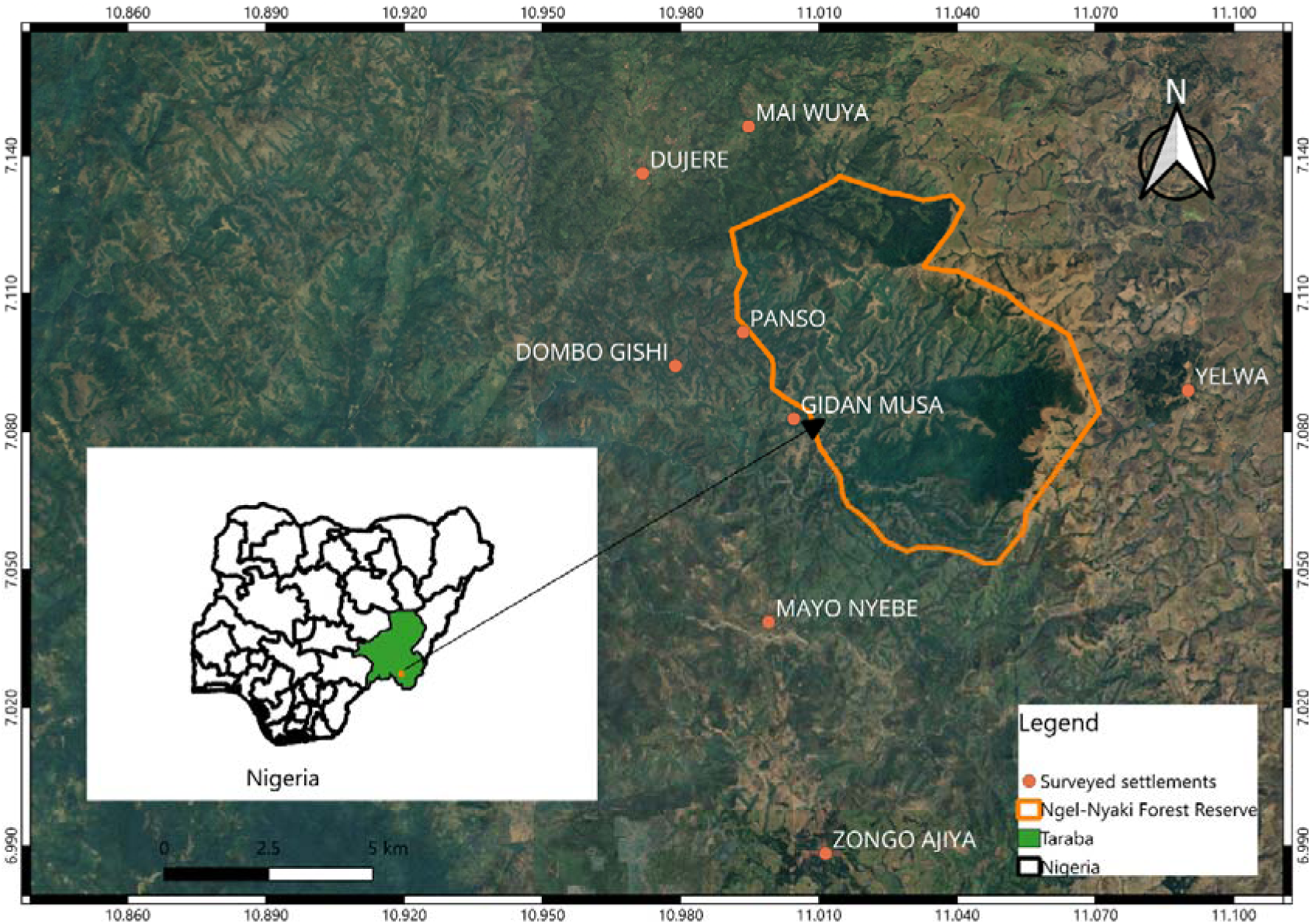
Study area showing the position of Ngel Nyaki Forest Reserve (green) and the settlements where interviews and focus group discussions were undertaken (orange-dots)

**TABLE 1.** Characteristics of the settlements included in the study.

| Name of Settlement | Elevation (m asl) | Presence of Eucalyptus Plantation | Distance to NNFR (Km) | # of Households | # of residents | Ethnic groups |
|---|---|---|---|---|---|---|
| Dombo Gishi | 976.72 | NO | 2 | 8 | 120 | Ndola |
| Dujure | 969.67 | NO | 1 | 86 | 900 | Mambilla, Ndola, Fulani |
| Gidan Musa | 1094.31 | NO | 0.5 | 5 | 55 | Ndola |
| Mai Wuya | 1130.54 | NO | 2 | 9 | 250 | Ndola, Fulani |
| Mayo Nyebe | 1076.29 | YES | 3 | 31 | 300 | Ndola, Fulani |
| Panso | 997.3 | NO | 1 | 7 | 50 | Manbilla, Ndola, Fulani |
| Yelwa | 1663.37 | YES | 3 | 216 | 2500 | Mambilla, Ndola, Kaka, Kambo, Fulani |
| Zongo Ajiya | 1370.88 | YES | 5 | 220 | 1500 | Mambilla, Ndola, Kaka, Fulani |

The ethnic composition varies considerably among the settlements. While most villages feature Mambilla or other Cameroonian borderland groups living alongside the Fulani, specific demographics range from homogeneous to highly mixed. For instance, Gidan Musa and Dombo Gishi are exclusively Ndola, whereas Panso, a settlement of comparable size, comprises Mambilla, Ndola, and Fulani residents. Yelwa is the most diverse settlement, containing speakers of all five ethnic languages. Overall, Ndola and Fulani are the commonly spoken languages across the study area.

### Data Collection

The study was undertaken between 2023 and 2024. We employed a participatory assessment (Cuni-Sanchez et al., 2019; Pingaronni et al., 2022) to gather community perceptions of the forest’s ES. This involved voluntary interviews with individuals and focus group discussions with residents from each of the eight settlements. Between 10 – 20 residents from each settlement were interviewed using the following questions:

1. Is Ngel-Nyaki Forest important for your community?
2. What benefits do you derive from the Ngel Nyaki Forest?
3. What plant species have benefitted you and your parents/grandparents?
4. What part/parts of the plant/tree species is/are used and how are they used?

For the focus group discussions in each settlement, our assessment included: i.) eliciting from residents a list of the perceived benefits (ES) derived from the forest reserve; ii.) ranking the ten most important services relevant to their settlements and iii.) identifying valuable plant species valued for provisioning services such as food, medicine, firewood, construction material, fodder, and honey. Honey was not grouped under food because it is an indirect service with its production depending on pollination, floral diversity and habitat quality; it is an outcome of pollination services rather than a primary provisioning service (Arany & Czúcz, 2025). The discussion groups included both male and female participants between the ages of 20 to 60 years old but were dominated by younger males (20-25 years old). We observed that the few women that were interviewed or were present at the discussions, almost always deferred to the men, even when they were asked to speak, explaining that they rarely went into the reserve but tended to stay closer to home. Interviews and discussions were carried out primarily in Fulfulde, given that it is now an almost-universal language in the area.

The ES identified by participants were grouped according to the types and sub-categories of the Millennium Ecosystem Assessment (Millennium Ecosystem Assessment, 2005). The plant species named in Fulfude, Mambilla or Ndola (Supplementary Table 3) were collected for translation and taxonomic identification by staff at the Nigerian Montane Forest Project’s herbarium.

### Data Analyses

Data on preferred ES by the local communities were compiled and compared across the settlements. Using the R statistical software version 4.3.1 (R Core Team, 2023), non-metric multidimensional scaling (NMDS) ordination was performed using the metaMDS function in the R package *vegan* (Oksanen et al., 2016) to examine the similarity in ecosystem perceptions across the settlements. To compare similarities in ecosystem service perceptions among settlements, we calculated the Jaccard similarity coefficient (*J*), which measures the proportion of shared ecosystem services between pairs of settlements based on presence–absence data (Jaccard, 1901). The index was calculated as:

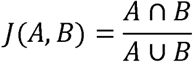

Where *A* and *B* represent the sets of ecosystem services identified in two different settlements, *A* ∩ *B* represents the number of ecosystem services shared between settlements, and *A* ⋃ *B* represents the total number of ecosystem services recognised across both settlements combined. Jaccard similarity values range from 0 to 1, where 1 indicates complete similarity in ecosystem service perceptions and 0 indicates complete dissimilarity.

### Ethics statement

At each settlement, we sought and received prior approval from the local community head called ‘Jauro’ to carry out the study. We also sought and received consent from community members before interviews and group discussions. We made sure they understood that participation was completely voluntary.

## RESULTS

### Ecosystem services identified by local communities

Twenty-four ES, comprising 12 provisioning, 6 regulating, 4 cultural, and 2 supporting services, were recorded across the eight settlements (Table 2). The number of ES identified per settlement ranged from 6 in Dujere (all provisioning) to 20 in Mayo Nyebe (Supplementary Table 1). The NMDS ordination (Figure 2) showed that provisioning services clustered near the centre of the ordination space which reflects that they were widely recognised by almost all the settlements. Food, honey and medicine were mentioned by all eight settlements (Supplementary Table 1 and Supplementary Figure 1). Fuelwood, timber, mushrooms, and lichens were each reported by seven settlements. Materials for roofing and crafts (e.g., basket making), alongside bushmeat, were cited in six settlements (Supplementary Figure 1). Despite this general overlap in provisioning services, some settlements reported some other ecosystem services. Mai Wuya, Gidan Musa and Yelwa clustered relatively closely together, indicating broadly similar perceptions of ecosystem services. Dombo Gishi and Dujure were separated from the other settlements along the negative side of the ordination plot. This suggests they recognised comparatively fewer ecosystem services. Services such as spiritual values, recreation, biodiversity support, water purification, microclimate regulation, and soil formation were positioned farther from the centre of the ordination, reflecting their less frequent mention across communities. Water provisioning and water flow regulation were reported in four settlements.

**FIGURE 2:**
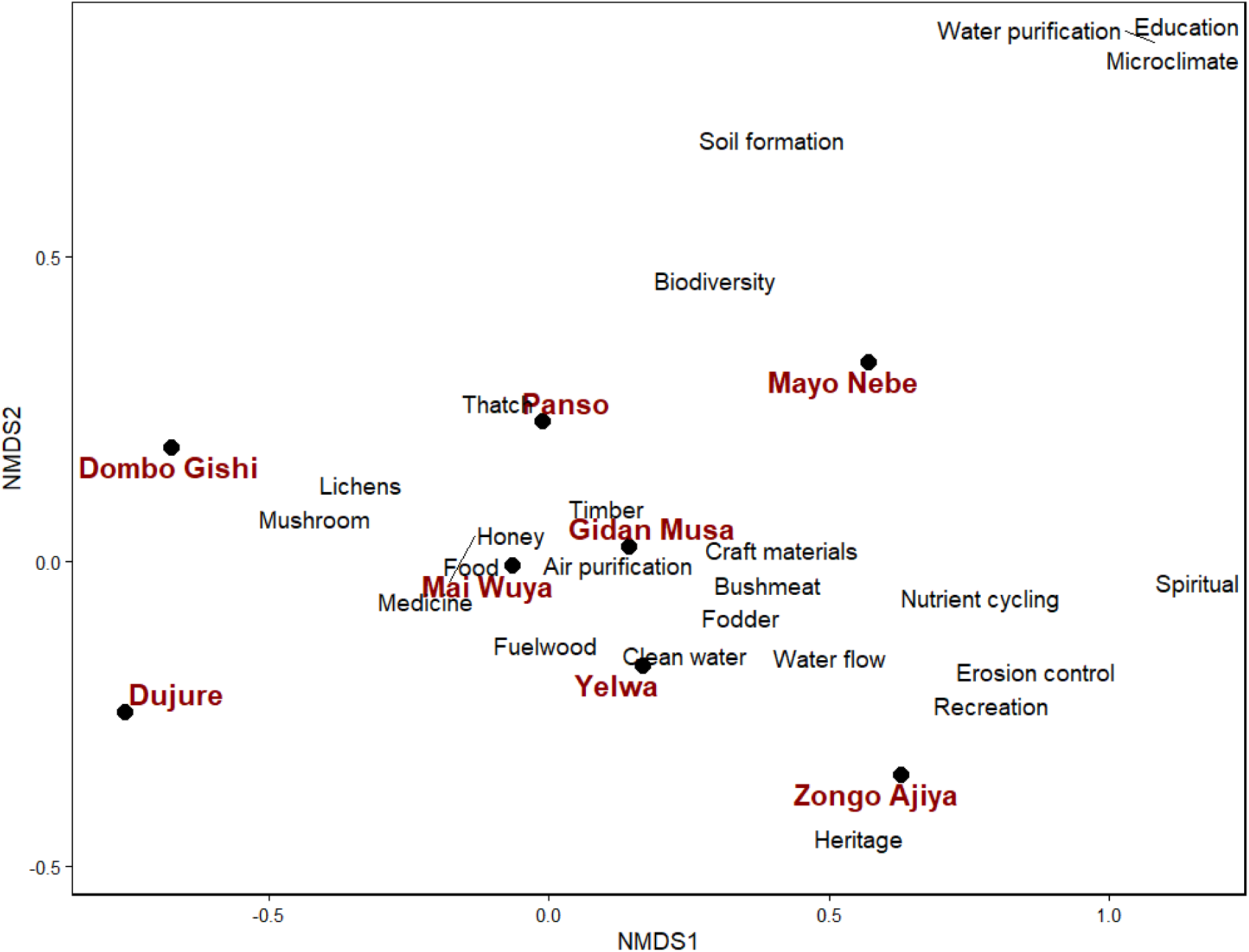
Non-metric multidimensional scaling (NMDS) ordination showing similarities in ecosystem services perception among eight settlements surrounding the Ngel Nyaki Forest Reserve (settlements positioned closer to each other had more similar perceptions on ecosystem services.

**Table 2.** Ecosystem services of Ngel Nyaki and surrounding grasslands as described by stakeholders from settlements.

| **Category** | **Services** |
|---|---|
| Provisioning Services | Food (vegetables, wild yams, fruits, nuts, fish, insects, wild fruits, condiments), honey, herbal medicine and poison, firewood, lichens, mushrooms, grass and dyes for thatch roofing and basket weaving, bush meat, water, fodder, timber, wood for crafting drums |
| Regulating Services | Water quantity and quality maintenance, air quality maintenance, Soil erosion control, nutrient cycling, microclimate regulation |
| Cultural Services | Heritage, sacred or religious sites, tourism and recreation site, education |
| Supporting Services | Biodiversity habitat, soil maintenance |

The Jaccard indices of similarity (J) ranged from 0.22 to 0.80, suggesting some variation in the degree to which settlements surrounding the reserve reported similar ecosystem services (Table 3). The highest similarity was observed between Mai Wuya and Yelwa (J = 0.80), followed by Mai Wuya and Panso (J = 0.73), and Gidan Musa and Panso (J = 0.72). In contrast there was least similarity between Dombo Gishi and Zongo Ajiya (J = 0.22) and between Dujure and Zongo Ajiya (J = 0.24). In general, Dombo Gishi and Dujere generally showed lower similarity with other settlements (Table 3).

**TABLE 3.** The Jaccard similarity indices in ecosystem services mentioned by the different settlements studied.

|  | **Dujure** | **Gidan Musa** | **Mai Wuya** | **Mayo Nebe** | **Panso** | **Yelwa** | **Zongo Ajiya** |
|---|---|---|---|---|---|---|---|
| **Dombo Gishi** | 0.62 | 0.41 | 0.58 | 0.35 | 0.50 | 0.47 | 0.22 |
| **Dujure** |  | 0.35 | 0.50 | 0.30 | 0.43 | 0.40 | 0.24 |
| **Gidan Musa** |  |  | 0.71 | 0.54 | 0.72 | 0.68 | 0.60 |
| **Mai Wuya** |  |  |  | 0.52 | 0.73 | 0.80 | 0.50 |
| **Mayo Nebe** |  |  |  |  | 0.55 | 0.59 | 0.52 |
| **Panso** |  |  |  |  |  | 0.61 | 0.45 |
| **Yelwa** |  |  |  |  |  |  | 0.67 |

### Communities ranking of most important Ecosystem Services

Table 4 presents the ES perceived by each community as providing the greatest benefits. Food was consistently ranked as the most important benefit across settlements, except for Mai Wuya where it was ranked second. Fuelwood provision was ranked first in the Mai Wuya settlement and appeared within the top five services for most other communities. In contrast, biodiversity support, recreation and tourism, and other cultural services (e.g. sacred groves and spiritual sites) were generally ranked in the lower half of the rankings.

**TABLE 4.** Ecosystem services ranking as perceived by different communities.

| Rank of benefits | Dombo Gishi | Dujure | Gidan Musa | Mai Wuya | Mayo Nyebe | Panso | Yelwa | Zongo Ajiya |
|---|---|---|---|---|---|---|---|---|
| 1 | Food | Food | Food | Fuelwood | Food | Food | Food | Food |
| 2 | Thatch | Fuelwood | Honey | Food | Thatch | Fuelwood | Honey | Medicines |
| 3 | Timber | Honey | Thatch | Honey | Medicines | Medicines | Lichens | Fuelwood |
| 4 | Medicines | Medicines | Fuelwood | Medicines | Fuelwood | Honey | Mushrooms | Timber |
| 5 | Honey | Mushrooms | Timber | Lichens | Timber | Water provisioning | Bushmeat | Thatch |
| 6 | Lichens | Lichens | Medicines | Thatch | Honey | Crafting materials | Fuelwood | Honey |
| 7 | Mushrooms |  | Biodiversity support | Timber | Crafting materials | Thatch | Medicines | Sacred grooves or spiritual sites |
| 8 |  |  | Water provisioning | Mushrooms | Sacred grooves or spiritual sites | Timber | Thatch | Crafting materials |
| 9 |  |  | Heritage ancestors-children | Water provisioning | Recreation and tourism | Bushmeat | Water provisioning | Recreation and tourism |
| 10 |  |  | Bush meat | Bush meat | Biodiversity support | Lichens | Timber | Fodder |

### Species identified by the communities for provisioning services

Overall, communities identified 97 plant species as providing provisioning ecosystem services (Supplementary Table 2). Of these, 30 species were cited as food sources (including spices and condiments), 44 species for medicinal purposes, 4 species for fuelwood, 6 species for construction purposes and 16 species for livestock fodder (Table 5). Several species were reported to have multiple uses.

**TABLE 5.**
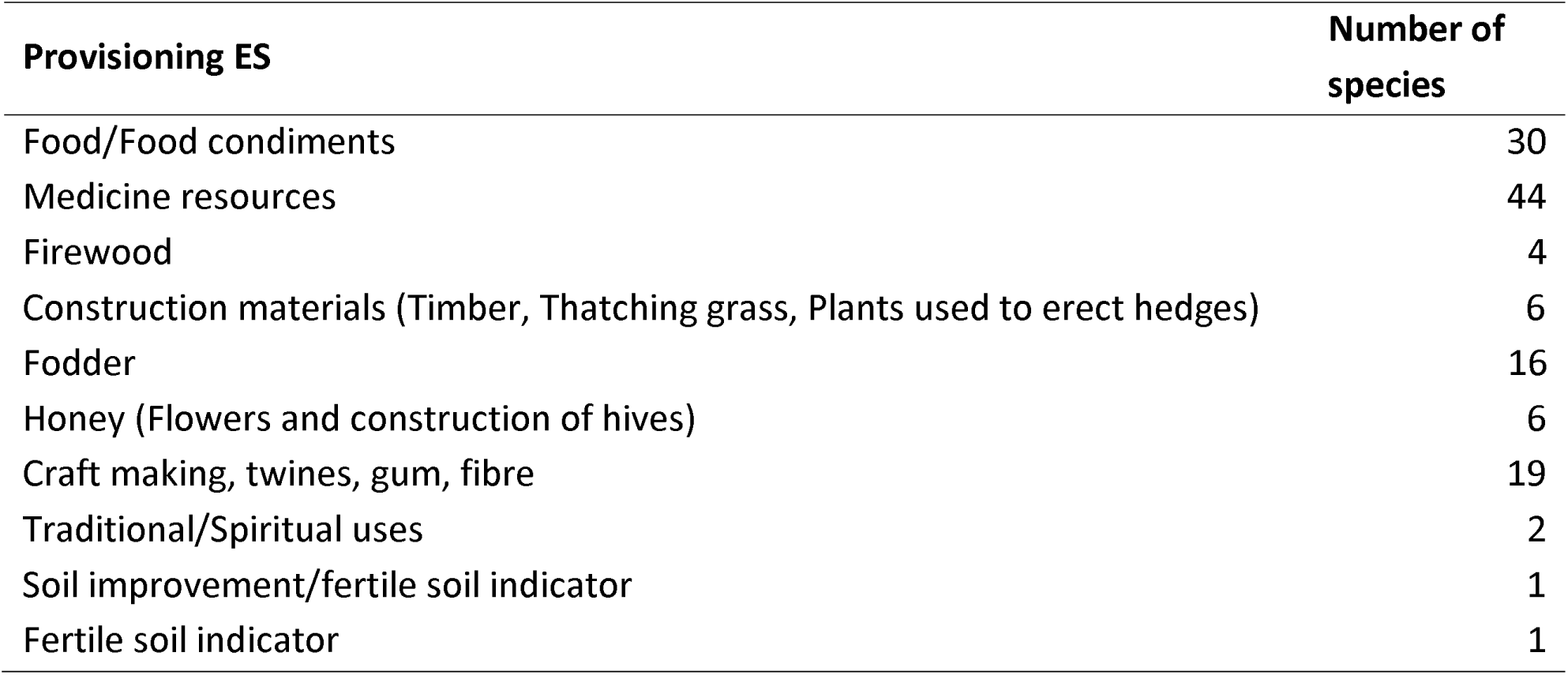
Total number of plant species reported by settlements for different provisioning services.

In terms of growth form, the recorded species comprised 62 trees, 15 shrubs, 6 climbers/lianas, 5 herbaceous perennials, 3 palms, 3 grasses, 1 fern, and 1 hemi-parasitic species (Supplementary Table 3).

Only five of the 97 documented species are classified as montane endemics. known to occur in the Ngel Nyaki forest. Thirty-five of the species are known to occur in the forest/forest edge, while the remaining species are characteristic of the surrounding savannah vegetation (Supplementary Table 3). Two of the recorded species—Chromolaena odorata and Azadirachta indica—are exotic species. Four of the species are listed in the IUCN Red Data List as Critically Endangered (Dombeya ledermannii and Eugenia gilgii), Endangered (Pterocarpus erinaceus) and Near Threatened (Polyscias fulva).

## DISCUSSION

Ngel Nyaki Forest Reserve is internationally valued for its exceptional Afromontane biodiversity (Key Biodiversity Areas Partnership, 2026). Our study, however, indicates that effective forest restoration requires balancing biodiversity conservation with the provision of ecosystem services valued by local stakeholders. Local communities identify provisioning services, derived from the nearby savannah and degraded grassland ecosystems more than the Afromontane forest, as the most valuable, underscoring the need to integrate these services into restoration planning.

### Perception of Ecosystem Services

The number of ES recognised by our interviewees varied three-fold among settlements, between 6 – 20. However, across all settlements provisioning services such as food (including honey), medicines and thatching materials were consistently ranked above regulating, cultural, or supporting services. This likely reflects the strong dependence these very poor and mostly uneducated rural communities have on natural resources for their livelihoods. It is a common pattern across tropical African highlands (e.g. Giday et al., 2016; Cuni-Sanchez et al. 2019; Fekadu et al., 2021; Miller et al., 2021) and sub-Saharan Africa more widely (Nyath et al., 2025; Ozcan et al., 2026). The universal recognition of honey as a provisioning service may, however, reflect the early efforts of Birdlife International and the Nigerian Conservation Foundation in promoting beekeeping through workshops in the area.

The finding that activities such as timber extraction and bush meat hunting, which are theoretically prohibited in the reserve, were nevertheless reported by some of the communities was somewhat unexpected. This contradicts the findings of Afriyie & Hecjmanová (2026), who reported that across Ghanaian protected areas supporting ES were perceived as the most important, with provisioning services less so because of strict no-access regulations. On the remote Mambilla plateau it is difficult to impose strict regulations; wealthy Fulani cattle owners can pay their way in to reserves and patrollers are under-paid and consequently unenthusiastic. This, combined with the fact that livelihoods are highly dependent on natural resources, highlights the need to strengthen initiatives that integrate conservation awareness with human wellbeing.

The lower value we found placed on regulating (e.g. water regulation, air purification, nutrient cycling) and supporting (e.g. biodiversity support, soil formation) services was not unexpected as indirect or less tangible ecosystem benefits are often less readily recognized by poor and uneducated populations (Zhang et al., 2016; Gouwakinnou et al., 2019; Wangchuk et al., 2021). While this likely reflects traditional knowledge (Khosravi Mashizi & Escobedo, 2025) and lower immediate utility, it highlights the undervaluation of services that underpin long-term ecosystem resilience. Of the settlements that did recognise regulating and supporting services, this was mostly in an indirect way. For instance, climate regulation, nutrient cycling and soil protection were mainly discussed in terms of how these help farms or health, not as separate ecological processes. For example, with comments such as “With the loss of the forests, the weather has become warmer and now there are more mosquitoes” and “the forest prevents heavy rains from destroying our farms”. Similar findings were reported by Berbés-Blázquez et al. (2016) and Comberti et al. (2015), demonstrating the need to use language, examples, and ideas that community members can relate to in conservation initiatives such as this (Gómez-Baggethun & Barton 2013).

Of note was that no one referred to crop pollination or pest control, even though several of the settlements are surrounded by subsistence farms. This was somewhat surprising given the importance of subsistence crops to local livelihoods, the strong recognition of honey as an ES, and the number of beekeepers among the respondents. Zhang et al. (2016) reported a similar unawareness of the link between pollination and crop yield for a more widespread study across rural Nigeria. This suggests that subsistence farmers need education on the link between their livelihood sources and the natural environment.

Cultural services were consistently ranked in the lower half of community priorities. This was unexpected given that Chapman & Chapman (2001), based on a 1970s study, reported the presence of sacred groves across the Mambilla Plateau that hold important spiritual and cultural significance for the Mambilla tribes. One possible explanation for this lack of recognition may be the high proportion of Fulani in our interviews-as a study in Benin (Gouwakinnou et al., 2019) found that Fulani communities generally exhibited low cultural affinities. In addition, the widespread influence of Christianity and Islam may have contributed to changes in traditional beliefs and cultural relationships with sacred natural sites. Future work should distinguish among Fulani, Christians and Muslims.

Alternatively, this apparent lack of cultural value may in reality reflect that the immediacy of livelihood needs overshadows the less tangible cultural benefits. In addition, cultural services are deeply embedded in traditions and belief systems that may not be readily articulated within a structured survey framework (Millennium Ecosystem Assessment, 2005; Daniel et al., 2012).

There is limited tourism-related activity in the area, however the fact that it was recognised suggests some awareness of the forest’s recreational potential, possibly through interactions with the Nigerian Montane Forest Project (NMFP) employees and researchers. The engagement of the NMFP with these communities has undoubtedly contributed to an understanding of the forest as a site of educational value.

### Variation in perception of ES across Settlements

The eight settlements included in this study were all within 5 km of the Ngel Nyaki forest edge but differed considerably in ethnic mix (Fulani, Ndola, Kaka, Mambilla, Kambu), population size (5 - 216 households), elevation (between 1600 – 900 m), distance to Afromontane/submontane forest (0.5 to 3 km) and proximity to Eucalyptus plantations (present or absent). Our sample size was insufficient for robust statistical modelling to determine the effect of these factors on settlement ES perceptions. However, simple observation of trends suggested none of these variables explained inter-settlement differences in ES perception. We found this surprising; e. g. the elevation gradient is extreme, and we had expected that respondents from the higher elevation settlements, especially Yelwa at 1600 m elevation, would have shown stronger affiliation with the montane/submontane species than settlements lower down. Another expectation was that settlements close to eucalyptus plantations would not perceive timber from the Reserve as an ES, because they would have access to eucalyptus. This was not the case as settlements with plantations, including Mayo Nyebe, Yelwa and Zongo Ajiya all mentioned timber in their 10 most important ecosystem services. This likely reflects the cost of Eucalyptus which has grown considerably over the years with the increase in its demand in lowland regions. Similarly, we had thought that the presence of a eucalyptus plantation would reduce the need for fuel wood, but apparently not. This may be because the owners of the eucalyptus plantations charge villagers excessively for leftover wood.

That distance to forest edge didn’t seem to influence response was also surprising. For example, Gouwakinnou et al. (2019) found that Benin communities closer to the forest edge identified more forest resources as important for their livelihoods and similar findings were reported for West Java (Muhamad et al., 2014). However, an alternative explanation may be that 5 km is close enough that all settlements used forest resources. There is very little agroforestry in the area, for example no orchards or fodder crops despite the suitable climate. This finding indicates that part of our restoration plan should involve a buffer zone with planted useful food and fodder plants (Damptey et al., 2021).

### Plant species for provisioning services

Of the 97 plant species identified by the communities as providing provisioning services, only 15 (16 %) were forest species and out of these, only five were true montane/submontane forest species (Abiem et al 2020). The remaining 84% of the species are common and widespread in the savannah. The forest species were mainly used for medicine and condiments, the savannah species provided a wide range of provisioning services including food, medicine and fodder. The savannah species fell into two groups, i) widespread and valued trees across the semi-arid Sahel such as Adansonia digitata, Afzelia africana, and Pterocarpus erinaceus (Assogbadjo et al., 2012; Nabaloum et al., 2022) and ii) widespread pioneer species such as Bridelia speciosa, Harungana madagascariensis and Trema orientalis associated with forest edges and disturbed areas (Abiem et al., 2020)). These pioneer species were also recorded by Cuni-Sanchez et al (2019) as being valued by highland communities in the Bamenda highlands of Cameroon.

Knowledge about these savannah/lowland species may have been introduced by the Fulani who migrated to the Mambilla Plateau in the early 20^th^ century and now settle on the Mambilla Plateau. They may have introduced and shared knowledge regarding the uses of these particular plant species with the local tribes on the Mambilla Plateau who now recognize them as valuable resources for their livelihood.

### Restoration implications for Ngel Nyaki Forest Reserve

When considering forest restoration and biodiversity, it is important to recognise the potential trade-offs between conservation goals and local communities’ preferences. Our findings suggest that the people living in close proximity to the reserve place greater value on the Savannah ecosystem than on the montane forest because the species in the savannah landscape provides them with many provisioning services that directly sustains their livelihoods. This suggests that restoration activities should consider the inputs of local communities which has been a major challenge in African restoration attempts (Bennett et al., 2009; Reed et al. 2016).

A central tension in ES-based restoration is balancing competing priorities: expanding forest cover can enhance carbon storage and other regulating services, and biodiversity but may reduce certain provisioning goods tied to savannah resources, while expanding savannah can boost provisioning goods but compromise biodiversity and carbon storage. This study’s findings underscore the practicality of trade-off analyses informed by local priorities. If provisioning ES overwhelmingly drive local well-being, immediate restoration actions that sustain or improve provisioning (e.g., replanting or protecting edible NTFP species, medicinal species, and food plants) can secure early community buy-in, which is critical for long-term success and enforcement of protected-area boundaries.

Our results argue for maintaining and expanding a mosaic landscape that includes both forest and derived savanna, rather than restoring the entire reserve into a closed-canopy forest. Such mosaic landscapes are characteristic of montane and sub-montane ecosystems across Africa where forest patches historically co-existed with savannah and grassland ecosystems (Fairhead & Leach, 1996). Such mosaics have historically mediated multiple ES, including provisioning (food, fodder, medicines) and regulating services (water storage, microclimate regulation, erosion control) (Tefera et al., 2024). Today, anthropogenic pressures-grazing, burning, and agricultural expansion—have shifted the balance toward grasslands in many areas, underscoring the need to explicitly account for landscape configuration in restoration planning.

The emphasis on a mosaic that includes derived savannah is particularly prescient. Derived (artificial) savannah ecosystems can offer substantial provisioning benefits (e.g., edible plants, fodder) (Osborne et al., 2018; Sankaran, 2019; Djiofack et al., 2024) and, depending on management practices, can contribute to carbon storage (Bastin et al., 2019; Zhou et al., 2023) and soil health (Teague & Kreuter, 2020). Importantly, the savannah component often remains crucial for livelihoods and cultural practices, and it can function as a tool for buffering biodiversity and ecological resilience when managed in conjunction with forest fragments. Restoration strategies should therefore not simply “add trees” but should aim to restore or maintain landscape heterogeneity and connectivity that support multiple ES pathways.

### Limitations of the study

Our study had several limitations with potential impacts on the interpretation of our findings. Firstly, we had a small sample size compared to similar studies that have been carried out in other mountain regions of Africa. Our study focused primarily on the major settlements located around the Ngel Nyaki Forest Reserve, and although it would have been desirable to include a larger number of settlements across the Mambilla Plateau, several communities were extremely remote and difficult to access. A further limitation is that the composition of interviewees and focus group participants may have also influenced the results. The Fulani tend to concentrate around larger settlements near the reserve. However, interviews and discussions did not distinguish between Fulani pastoralists and subsistence farming communities. As a result, potentially important differences in ecological knowledge, livelihood dependence, and ecosystem service preferences between these groups may not have been captured. Future studies would benefit from explicitly separating these stakeholder groups to better understand how cultural background, livelihood strategies, and mobility influence ecosystem service valuation.

Finally, although the study provides important insights into local perceptions surrounding the only formally protected montane forest reserve on the Mambilla Plateau, a wider resurvey incorporating additional settlements and a more standardised sampling framework would improve understanding of ecosystem service priorities across the broader Plateau landscape.

## CONCLUSIONS AND RECOMMENDATIONS

In summary, our study highlights that effective restoration of Ngel Nyaki Forest Reserve must explicitly balance conservation goals with the ecosystem services people depend on. This calls for integrated restoration planning that blends forest recovery with community needs. For example, we recommend establishing buffer zones planted with fruit, fodder, and medicinal plants valued by locals, expanding community-driven initiatives such as sustainable beekeeping, and involving stakeholders in co-management of forest resources. Policymakers should adapt reserve regulations to allow carefully regulated access to non-timber forest products and support participatory management, aligning conservation with livelihood security. Further research and monitoring are also needed to evaluate how these combined strategies affect both biodiversity and human well-being. By openly addressing these trade-offs – recognizing that strict protection alone may not meet local priorities – an adaptive, community-centred restoration approach may ultimately safeguard the reserve’s exceptional biodiversity while also improving local livelihoods.

## ACKNOWLEDGEMENTS

The authors are exceedingly grateful to the Community heads ‘Jauros’ and people of Dombo Gishi, Dujere, Gidan Musa, Mai Wuya, Panso, Mayo Yembe, Yelwa and Zongo Ajiya communities for giving us permission to carry out this study and for their time and enthusiasm in sharing their knowledge with us. We also wish to thank the management and staff of the Nigerian Montane Forest Project who provided accommodation, field assistance and translation services during field work.

