## Supplementary tables for "Indigenous communities’ perceptions of ecosystem services of a montane forest ecosystem in Nigeria: Implications for restoration planning"

**SUPPLEMENTARY INFORMATION**

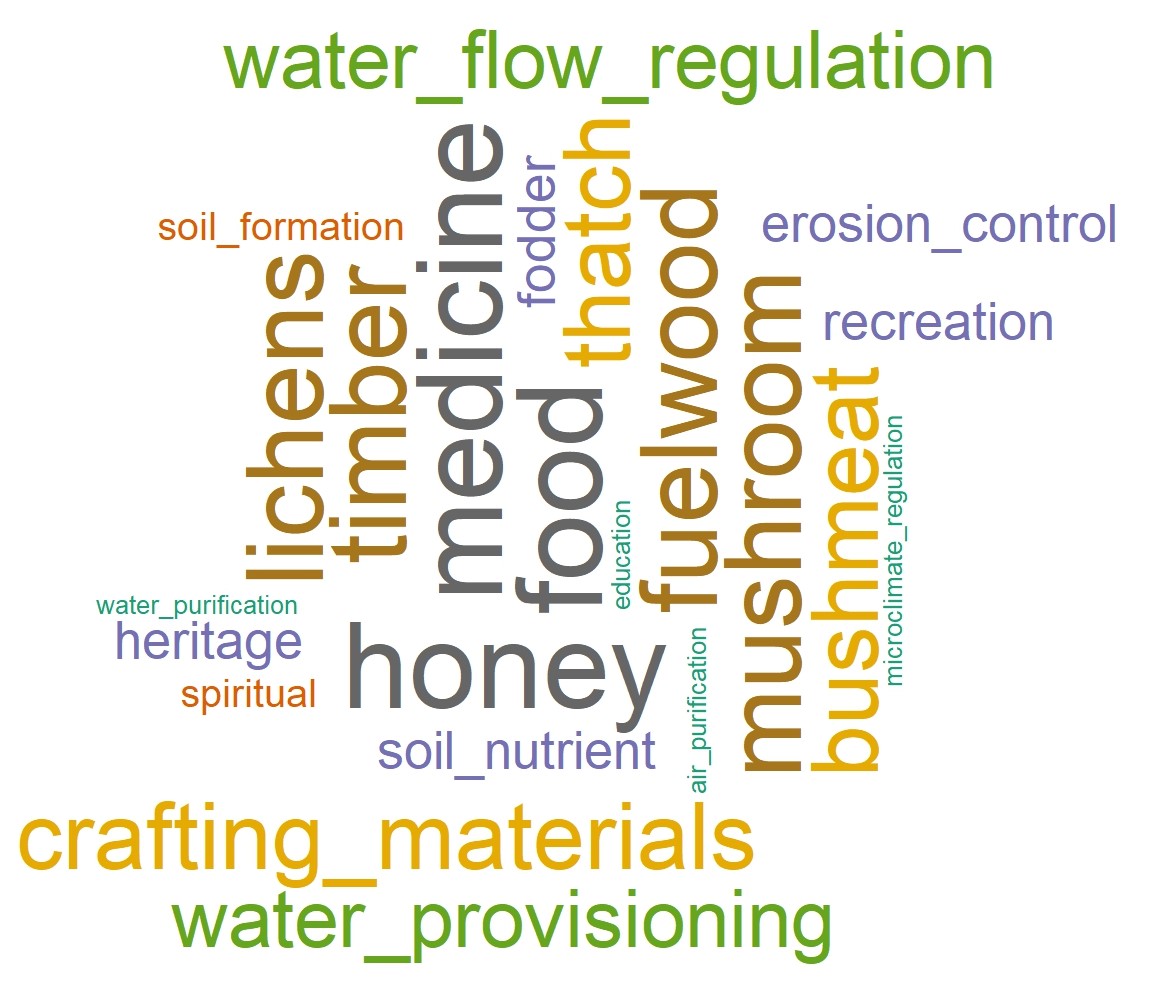

**Supplementary Figure 1: A cloud of participant responses on Ecosystem Services (font size is directly proportional to the frequency of the words occurrence)**

**Supplementary Table 1: Number of ecosystem services recognised in the focus-group discussions (FGDs) by community**

|  | | **Dombo Gishi** | **Dujure** | **Gidan Musa** | **Mai Wuya** | **Mayo Nyebe** | **Panso** | **Yelwa** | **Zongo Ajiya** |
| --- | --- | --- | --- | --- | --- | --- | --- | --- | --- |
| **No. ES mentioned** | | 7 | 6 | 17 | 12 | 20 | 14 | 15 | 15 |
| Provisioning | Food (vegetables, wild yams, fruits, nuts, fish, insects) | * | * | * | * | * | * | * | * |
|  | Honey | * | * | * | * | * | * | * | * |
|  | Medicines | * | * | * | * | * | * | * | * |
|  | Fuelwood |  | * | * | * | * | * | * | * |
|  | Timber | * |  | * | * | * | * | * | * |
|  | Mushroom | * | * | * | * | * | * | * |  |
|  | Bush meat |  |  | * | * | * | * | * | * |
|  | Crafting materials |  |  | * | * | * | * | * | * |
|  | Thatching materials | * |  | * | * | * | * | * |  |
|  | Fodder |  |  | * |  |  | * |  | * |
|  | Lichens | * | * | * | * | * | * | * |  |
|  | Clean water |  |  | * | * |  | * | * | * |
| Regulating | Microclimate regulation |  |  |  |  | * |  |  |  |
|  | Nutrient cycling |  |  | * |  | * |  |  | * |
|  | Erosion control |  |  |  |  | * |  | * | * |
|  | Water flow regulation |  |  | * | * | * |  | * | * |
|  | Water purification |  |  |  |  | * |  |  |  |
|  | Air purification |  |  | * |  |  |  |  |  |
| Cultural | Heritage |  |  | * |  |  |  | * | * |
|  | Sacred groves/ spiritual sites |  |  |  |  | * |  |  | * |
|  | Recreation/tourism |  |  |  |  | * |  | * | * |
|  | Education |  |  |  |  | * |  |  |  |
| Supporting | Biodiversity support |  |  | * |  | * | * |  |  |
|  | Soil formation |  |  |  |  | * | * |  |  |

**Supplementary Table 2: Plant species identified by the communities that are used for provisioning services**

| **Species** | **Family** | **Habit** | **Habitat** | **Montane Endemic?** | **Note on use** |
| --- | --- | --- | --- | --- | --- |
| *Afzelia africana* Sm. ex Pers. | Fabaceae | Tree | Savannah woodland | No | Animal fodder |
| *Albizia* sp. | Fabaceae | Tree | Savannah woodland | No | Making fishing tool |
| *Alchornea laxiflora* (Benth.) Pax & K.Hoffm. | Phyllanthaceae | Tree/Shrub | Savannah woodland | No | Leaves used for medicine |
| *Aframomum melegueta* K.Schum. | Zingiberaceae | Herbaceous perennial | Forest Edge /Savannah | No | Seeds used as condiment |
| *Allophylus africanus* P.Beauv. | Sapindaceae | Small tree/shrub | Forest | No | Leaves used as fodder and gum |
| *Adansonia digitata* L. | Malvaceae | Tree | Savannah | No | Fruits and leaves are eaten |
| *Annona senegalensis* Pers. | Annonaceae | Small tree/shrub | Savannah | No | Fruit edible and leaves used for medicine |
| *Anogeissus leiocarpa* (DC.) Guill. & Perr. | Combretaceae | Tree | Savannah | No | Animal fodder |
| *Anonychium africanum* (Guill. & Perr.) C.E.Hughes & G.P.Lewis | Fabaceae | Tree | Savannah | No | Used as timber |
| *Anthocleista vogelii* Planch. | Gentianaceae | Tree | Forest | No | Bark used for medicine |
| *Anthonotha noldeae* A.Chev. | Fabaceae | Tree | Forest | Yes | Bark used for medicine |
| *Aspilia africana* (Pers.) C.D.Adams | Asteraceae | Semi-woody herb | Savannah | No | Animal fodder |
| *Azadirachta indica* A.Juss. | Meliaceae | Tree | Exotic species | No | Used for medicine |
| *Beilschmiedia mannii* (Meisn.) Benth. & Hook.f. | Lauraceae | Tree | Forest | No | Fruit valuable food and leaves used as condiment |
| *Bombax costatum* Pellegr. & Vuillet | Malvaceae | Tree | Savannah | No | Silk used for textile; dry flower used as condiment |
| *Borassus aethiopum* Mart. | Arecaceae | Palm | Savannah | No | Fruits, seeds and sprouts of germinated seeds are eaten |
| *Bridelia ferruginea* Benth. | Phyllanthaceae | Tree/Shrub | Forest/ Savannah | No | Animal fodder |
| *Bridelia* sp. | Phyllanthaceae | Tree/Shrub | Forest/ Savannah | No | Animal fodder, Honey |
| *Calotropis procera* (Aiton) Dryand. | Apocynaceae | Shrub | Savannah | No | Used for medicine |
| *Campylospermum flavum* (Vahl) Farron | Ochnaceae | Small tree/shrub | Forest | No | Bark used for medicine |
| *Canthium* sp. | Rubiaceae | Small tree/shrub | Forest | No | Spiritual significance |
| *Carapa oreophila* Kenfack | Meliaceae | Tree | Forest | Yes | Fruit crushed and fed to animals |
| *Chromolaena odorata* (L.) R.M.King & H.Rob. | Asteraceae | Perennial herb | savannah | No (Invasive) | Used as indicator for fertile land |
| *Clematis hirsuta* Guill. & Perr. | Ranunculaceae | Perennial climber | Forest/ Savannah | No | Used for medicine |
| *Combretum molle* R.Br. ex G.Don | Combretaceae | Tree | Savannah | No | Fuelwood |
| *Crossopteryx febrifuga* (Afzel. ex G.Don) Benth. | Rubiaceae | Small tree/shrub | Savannah | No | Fuelwood, Honey and medicine |
| *Croton macrostachyus* Hochst. ex Delile | Euphorbiaceae | Tree | Forest Edge/ Savannah | No | Fuelwood and medicine |
| *Cussonia arborea* Hochst. ex A.Rich. | Araliaceae | Tree | Savannah | No | Used for making farming tools and other crafts |
| *Daniellia oliveri* (Rolfe) Hutch. & Dalziel | Fabaceae | Tree | Savannah | No | Leaves used as fodder; gum is extracted from trunk |
| *Deinbollia* sp. | Sapindaceae | Tree | Forest | No | Leaves used for medicine |
| *Dichrostachys cinerea* (L.) Wight & Arn. | Fabaceae | Shrub | Savannah | No | Fruits used for fodder; bark used for medicine |
| *Dioscorea* sp. | Dioscoreaceae | Climber | Forest Edge/ Savannah | No | Wild yam used for food |
| *Dissotis* sp. | Melastomataceae | Herbaceous perennial | Savannah | No | Fresh leaves used for medicine |
| *Dombeya ledermannii* Engl. | Malvaceae | Tree | Forest Edge/ Savannah | Yes | Fibre used for twines, Honey |
| *Elaeis guineensis* Jacq. | Arecaceae | Palm | Planted | No | Fruits used for several purposes |
| *Entada abyssinica* A.Rich. | Fabaceae | Small tree/shrub | Savannah | No | Fruits and leaves used for fodder |
| *Erythrina senegalensis* DC. | Fabaceae | Small tree/shrub | Savannah | No | Bark and seed used for medicine; leaves used for fodder |
| *Erythrina sigmoidea* Hua | Fabaceae | Small tree/shrub | Savannah | No | Bark and seed used for medicine; leaves used for fodder |
| *Eugenia gilgii* Engl. & Brehmer | Myrtaceae | Small tree/shrub | Forest | No | Fruits are eaten and leaves are used for medicine |
| *Ficus* sp. | Moraceae | Tree | Forest/ Savannah | No | Used for erecting hedges and for medicine |
| *Ficus sur* Forssk. | Moraceae | Tree | Forest/ Savannah | No | Used for erecting hedges and for medicine |
| *Garcinia* sp. | Clusiaceae | Tree | Forest | No | Used for medicine |
| *Gardenia erubescens* Stapf & Hutch. | Rubiaceae | Shrub | Savannah | No | Fruits are eaten, wood used by blacksmith |
| *Harungana madagascariensis* Lam. ex Poir. | Hypericaceae | Small tree/shrub | Savannah | No | Used to craft utensils |
| *Hyparrhenia rufa* (Nees) Stapf | Poaceae | Grass | Savannah | No | Used for thatched roofing |
| *Hymenocardia acida* Tul. | Phyllanthaceae | Small tree/shrub | Savannah | No | Leaves, bark and roots used for medicine |
| *Ilex mitis* (L.) Radlk. | Aquifoliaceae | Small tree/shrub | Savannah | No | Used for fodder |
| *Imperata cylindrica* (L.) Raeusch. | Poaceae | Grass | Savannah | No | Used for craft |
| *Jatropha curcas* L. | Euphorbiaceae | Small tree/shrub | Savannah | No | Leaves and seeds used for medicine |
| *Khaya grandifoliola* C.DC. | Meliaceae | Tree | Forest | No | Used for medicine and for timber |
| *Khaya senegalensis* (Desr.) A.Juss. | Meliaceae | Tree | Savannah | No | Bark, Seed oil and gum used for medicine |
| *Kigelia africana* (Lam.) Benth. | Bignoniaceae | Tree | Forest/Savannah | No | Bark and fruits used for medicine |
| *Landolfia* sp. (Small fruit) | Apocynaceae | Liana | Forest/Savannah | No | Fruits are eaten, fibre used for twine |
| *Landolfia* sp. (large fruit) | Apocynaceae | Liana | Forest/Savannah | No | Fruits are eaten, fibre used for twine |
| *Lannea* sp. | Anacardiaceae | Tree | Savannah | No | Bark used for medicine |
| Unidentified liana | Unknown | Liana | Forest | No | Used for twine |
| *Lophira lanceolata* Tiegh. ex Keay | Ochnaceae | Tree | Savannah | No | Shoot used for medicine |
| *Maesa lanceolata* Forssk. | Primulaceae | Small tree/shrub | Forest/Savannah | No | Used to craft hunting tools |
| Mistletoe | Loranthaceae | Hemi-Parasite | Forest/Savannah | No | Used for medicine |
| *Mussaenda arcuata* Poir. | Rubiaceae | Liana | Forest/Savannah | No | Fruits are eaten |
| *Ocimum* sp. | Lamiaceae | Shrub | Savannah | No | Leaved are used as a spice |
| *Parkia biglobosa* (Jacq.) R.Br. ex G.Don | Fabaceae | Tree | Savannah | No | Fruit pulp is eaten and used for medicine; seeds are used to make condiment |
| *Phoenix reclinata* Jacq. | Arecaceae | Palm | Savannah | No | Fruit edible and palm fronds use in craft making |
| *Phyllanthus muellerianus* (Kuntze) Exell | Phyllanthaceae | Shrub | Savannah | No | Used for food and for medicine |
| *Piliostigma thonningii* (Schumach.) Milne-Redh. | Fabaceae | Small tree/shrub | Savannah | No | Used for animal fodder and used to make craft |
| *Piper* sp. | Piperaceae | Shrub | Forest | No | Leaves are eaten and seeds used as spice |
| *Pittosporum viridiflorum* Sims | Pittosporaceae | Tree | Forest Edge/Savannah | No | fruits and seeds are used as spice |
| *Polyscias fulva* (Hiern) Harms | Araliaceae | Tree | Forest | No | Trunk used for making traditional drums and for medicine |
| *Protea madiensis* Oliv. | Proteaceae | Shrub | Savannah | No | Roots used for medicine |
| *Prunus africana* (Hook.f.) Kalkman | Rosaceae | Tree | Forest Edge/Savannah | Yes | Bark used for medicine |
| *Pteridium aquilinum* (L.) Kuhn | Dennstaedtiaceae | Fern | Savannah | No | Weed and sometimes fibre use for twine |
| *Pterocarpus erinaceus* Poir. | Fabaceae | Tree | Savannah | No | Timber; Bark used for medicine |
| *Rauvolfia vomitoria* Afzel. | Apocynaceae | Small tree/shrub | Forest | No | Bark used for medicine |
| *Ricinus communis* L. | Euphorbiaceae | Shrub | Savannah | No | Seed used for producing oil used for medicine |
| *Ricinodendron heudelotii* (Baill.) Heckel | Euphorbiaceae | Tree | Savannah | No | Trunk used for making Drum |
| *Rubus* sp. | Rosaceae | Shrub | Savannah | No | Fruits are eaten |
| *Rumex abyssinicus* Jacq. | Polygonaceae | Herbaceous perennial | Savannah | Yes | Used as condiment |
| *Nauclea latifolia* Sm. | Rubiaceae | Small tree/shrub | Savannah | No | Roots used for medicine |
| *Senna* sp. | Fabaceae | Tree | Savannah | No | Used for animal fodder |
| *Sida acuta* Burm.f. | Malvaceae | Subshrub | Savannah | No | Used for medicine |
| *Solanum* sp. | Solanaceae | Shrub | Savannah | No | Seeds used for traditional and religious purpose |
| *Sporobolus pyramidalis* P.Beauv. | Poaceae | Grass | Savannah | No | Young shoots used as fodder and mature grass used in craft making |
| *Symphonia globulifera* L.f. | Clusiaceae | Tree | Forest | No | Bark used for medicine |
| *Syzygium macrocarpum* Engl. | Myrtaceae | Tree | Savannah | No | Fruits are eaten; leaves are used for medicine, Honey |
| *Syzygium guineense* (Willd.) DC. | Myrtaceae | Tree | Forest Edge/Savannah | No | Fruits are eaten; leaves are used for medicine |
| *Tamarindus indica* L. | Fabaceae | Tree | Savannah | No | Fruits are used as condiment |
| *Tephrosia vogelii* Hook.f. | Fabaceae | Shrub | Savannah | No | Plant used on farmlands to improve soil fertility and for medicine |
| *Terminalia ivorensis* A.Chev. | Combretaceae | Tree | Savannah | No | Wood used for fuel wood |
| *Trema orientalis* (L.) Blume | Cannabaceae | Small tree/shrub | Forest Edge/ Savannah | No | Leaves used for medicine |
| *Uapaca togoensis* Pax | Phyllanthaceae | Tree | Savannah | No | Fruits are eaten, used to set up beehives |
| *Uvaria chamae* P.Beauv. | Annonaceae | Shrub | Savannah | No | Fruits are eaten |
| *Vepris* sp. | Rutaceae | Shrub | Forest Edge/ Savannah | No | Leaves used for medicine |
| *Vernonia amygdalina* Delile | Asteraceae | Shrub | Savannah | No | Leaves are used for food and for medicine |
| *Vitex doniana* Sweet | Lamiaceae | Tree | Savannah | No | Fruits are eaten and wood is good timber, Honey |
| *Xylopia* sp. | Annonaceae | Small tree/shrub | Savannah | No | Fruits are used as spice |
| *Zanthoxylum* sp. | Rutaceae | Tree | Savannah | No | Fruits are used as condiments; roots for medicine |

| **Supplementary Table 3: Local names of plant species used for different provisioning ecosystem services** | | | |
| --- | --- | --- | --- |
| **Species** | **Local Names** | | |
|  | **Mambilla** | **Fulfude** | **Ndola** |
| *Afzelia africana* Sm. ex Pers. |  | Kawohi |  |
| *Albizia* sp. | Yomen | Yamerehi |  |
| *Alchornea laxiflora* (Benth.) Pax & K.Hoffm. |  | Karkandam |  |
| *Aframomum melegueta* K.Schum. | Gwa | Sitta lainde |  |
| *Allophylus africanus* P.Beauv. | Tager | Buduchi |  |
| *Adansonia digitata* L. |  | Bokki |  |
| *Annona senegalensis* Pers. |  | Dukkuhi | Aboha |
| *Anogeissus leiocarpa* (DC.) Guill. & Perr. |  | Gosgahi |  |
| *Anonychium africanum* (Guill. & Perr.) C.E.Hughes & G.P.Lewis |  |  |  |
| *Anthocleista vogelii* Planch. | Taroo |  |  |
| *Anthonotha noldeae* A.Chev. |  | Kojoli |  |
| *Aspilia africana* (Pers.) C.D.Adams | We-el | Sonyonai |  |
| *Azadirachta indica* A.Juss. |  | Neem |  |
| *Beilschmiedia mannii* (Meisn.) Benth. & Hook.f. | Ngwe | Konkoli |  |
| *Bombax costatum* Pellegr. & Vuillet | Zankum | Bantahi | Pumma |
| *Borassus aethiopum* Mart. |  | Dub-i |  |
| *Bridelia ferruginea* Benth. |  | Bududi dimi | Akwaa |
| *Bridelia* sp. | Jar | Mburumburum |  |
| *Calotropis procera* (Aiton) Dryand. |  |  |  |
| *Campylospermum flavum* (Vahl) Farron |  | Kokarahi |  |
| *Canthium* sp. |  | Jagarahi |  |
| *Carapa oreophila* Kenfack |  | Karangahi/ Karangatu |  |
| *Chromolaena odorata* (L.) R.M.King & H.Rob. |  | Bukasa |  |
| *Clematis hirsuta* Guill. & Perr. |  |  |  |
| *Combretum molle* R.Br. ex G.Don | Veer | Sakatahi |  |
| *Crossopteryx febrifuga* (Afzel. ex G.Don) Benth. |  | Remajogohi |  |
| *Croton macrostachyus* Hochst. ex Delile | Buor | Ngalawahi |  |
| *Cussonia arborea* Hochst. ex A.Rich. | Banker | Hoyahi | Adop |
| *Daniellia oliveri* (Rolfe) Hutch. & Dalziel |  | Kayalahi |  |
| *Deinbollia* sp. | Gib |  |  |
| *Dichrostachys cinerea* (L.) Wight & Arn. | Tuo njerre | Burle |  |
| *Dioscorea* sp. |  |  |  |
| *Dissotis* sp. | Ndakal | Gaku |  |
| *Dombeya ledermannii* Engl. | Te-el | Dalamhi |  |
| *Elaeis guineensis* Jacq. | Mba'a | Kwaraje |  |
| *Entada abyssinica* A.Rich. | Bo'om | Pelwahi |  |
| *Erythrina senegalensis* DC. | Gomm | Bordilohi | Ndeya |
| *Erythrina sigmoidea* Hua | Gomm | Bantahi | Ndeya |
| *Eugenia gilgii* Engl. & Brehmer |  | Muramoda |  |
| *Ficus* sp. | Boll | Cediyahi |  |
| *Ficus sur* Forssk. | Tagar | Ibbe |  |
| *Garcinia* sp. | Nger | Dankamaru | Ambata |
| *Gardenia erubescens* Stapf & Hutch. | Ndeng | Dingali |  |
| *Harungana madagascariensis* Lam. ex Poir. | Jeou | Burugalhi |  |
| *Hyparrhenia rufa* (Nees) Stapf |  |  |  |
| *Hymenocardia acida* Tul. | Korr | Yewasettohi |  |
| *Ilex mitis* (L.) Radlk. |  | Ndanehohi |  |
| *Imperata cylindrica* (L.) Raeusch. | Liel | Sojo |  |
| *Jatropha curcas* L. |  | Kwalakwalahi |  |
| *Khaya grandifoliola* C.DC. |  |  |  |
| *Khaya senegalensis* (Desr.) A.Juss. |  | Kahi |  |
| *Kigelia africana* (Lam.) Benth. |  | Jirlahi |  |
| *Landolfia* sp. (Small fruit) | Larum | Tibokko |  |
| *Landolfia* sp. (large fruit) | Mamber | Pogije |  |
| *Lannea* sp. |  | Sonyi | Aruma |
| Unidentified liana | Lir | Lai |  |
| *Lophira lanceolata* Tiegh. ex Keay | Tuo nu | Karehi |  |
| *Maesa lanceolata* Forssk. |  | Belbelhi |  |
| Mistletoe | Dalap | Sohore |  |
| *Mussaenda arcuata* Poir. |  | Bue pabi |  |
| *Ocimum* sp. |  |  |  |
| *Parkia biglobosa* (Jacq.) R.Br. ex G.Don | Manyer | Narehi |  |
| *Phoenix reclinata* Jacq. | Zorr | Dalle |  |
| *Phyllanthus muellerianus* (Kuntze) Exell |  | Ribribi | Simsa |
| *Piliostigma thonningii* (Schumach.) Milne-Redh. |  | Barkehi | Bartejo |
| *Piper* sp. | Lomm |  |  |
| *Pittosporum viridiflorum* Sims |  | Nyanyulehi |  |
| *Polyscias fulva* (Hiern) Harms | Ruon | Lekko mbaggu |  |
| *Protea madiensis* Oliv. |  | Lebrehi |  |
| *Prunus africana* (Hook.f.) Kalkman |  |  |  |
| *Pteridium aquilinum* (L.) Kuhn | Enkon | Agugu |  |
| *Pterocarpus erinaceus* Poir. | Tuo homme | Banohi | Ajengha |
| *Rauvolfia vomitoria* Afzel. |  | Kofa |  |
| *Ricinus communis* L. | Chur | Jurman | Angeren |
| *Ricinodendron heudelotii* (Baill.) Heckel |  |  | Kuma |
| *Rubus* sp. | Tambul borre | Nyamdu baka |  |
| *Rumex abyssinicus* Jacq. |  |  |  |
| *Nauclea latifolia* Sm. |  | Bakurehi | Afula |
| *Senna* sp. |  |  |  |
| *Sida acuta* Burm.f. | Weé | Saldori | Nyinna |
| *Solanum* sp. | Ngoó | Kuwitaje |  |
| *Sporobolus pyramidalis* P.Beauv. | Gwur | Pagame |  |
| *Symphonia globulifera* L.f. |  | Kandihi |  |
| *Syzygium macrocarpum* Engl. | Enkan | Malmoje |  |
| *Syzygium guineense* (Willd.) DC. | Tawur | Sumsum |  |
| *Tamarindus indica* L. |  | Jab-i |  |
| *Tephrosia vogelii* Hook.f. | Twin | Yomji |  |
| *Terminalia ivorensis* A.Chev. | Thor | Kulahi | Attaa |
| *Trema orientalis* (L.) Blume |  | Gauri chollihi |  |
| *Uapaca togoensis* Pax |  |  |  |
| *Uvaria chamae* P.Beauv. |  |  |  |
| *Vepris* sp. |  | Komhi |  |
| *Vernonia amygdalina* Delile | Luwe | Suwankali |  |
| *Vitex doniana* Sweet | Tergen | Bummeje | Galbije |
| *Xylopia* sp. |  | Kimbhi |  |
| *Zanthoxylum* sp. |  | Pasakorihi |  |
